# Effects of healthy ageing on activation pattern within the brain during movement and motor imagery: an fMRI study

**DOI:** 10.1101/584813

**Authors:** Sanjay Budhdeo, Jean-Claude Baron, Nikhil Sharma

## Abstract

**Introduction:** Motor imagery (MI) has potential as an intervention to improve performance in neurological disease affecting the motor system and to modulate brain computer interfaces (BCI). We hypothesized that the shared networks of MI and executed movement (EM) would be affected by age. Understanding these changes is important in application of MI in neurological disorders.

**Methods:** Using tensor-independent component analysis (TICA), we mapped the neural networks involved during MI and EM in 31 healthy volunteers (ages 20-72), who were recruited and screened for their ability to perform imagery. We used an fMRI block-design with MI & rest and EM & rest.

**Results:** TICA defined 37 independent components (ICs). Eight remained after excluding ICs representing artifacts. These ICs accounted for 35% of variance. While all ICs had greater activation in EM than MI. Two ICs increased with greater age for EM only. These ICs contained a bilateral network of brain areas, including primary motor cortex and cerebellum.

**Conclusion:** This study demonstrates the prominence of shared cerebral networks between MI and EM. There are age-dependent changes to EM activation, while MI activation appeared age independent. This strengthens the rationale for using MI to access the motor networks independent of age.

## Introduction

Neurological diseases affecting the motor system are increasing in incidence with an ageing population (Pakpoor & Goldacre, 2017). Two important drivers of this increase are neurodegenerative disease, including motor neuron disease, and brain injury such as stroke. Many neurodegenerative diseases result in functional limitations in mobility and movement. Motor imagery has potential as an intervention to improve motor performance and can be used to drive BCI (Sharma, Pomeroy, & Baron, 2006). In this study, we examined the networks shared between motor imagery (MI) and executed movement (EM) with age.

Motor imagery can be defined as a dynamic state during which the representation of a specific motor action is internally reactivated within working memory without any overt motor output, and that is governed by the principles of central motor control (Sharma et al., 2006). Motor imagery provides an alternative means to access the motor system, that in principle, is not influenced by age related changes outside of the CNS. Motor imagery represents a potential method to access the motor system to encourage to improve performance in patients with neurological diseases.

There are several reasons for considering a relationship between motor imagery and execution. First, there is close temporal coupling between motor imagery and movement (Decety, Jeannerod, & Prablanc, 1989). During imagined movement, the reduction in accuracy with increasing speed (i.e. Fitt’s Law) is maintained (Decety & Jeannerod, 1995). Second, there is cross-modality evidence of similarities between brain activity from motor imagery and motor execution. Electrophysiological studies suggest that in both cases, mu topographies showed bilateral foci of desynchronization over sensorimotor cortices, while beta topographies showed peak desynchronization over the vertex. Multivariate fMRI analysis that we have undertaken demonstrates that the majority of the networks involved in motor imagery and execution appear to be shared.

We have previously reported that motor imagery and execution involves a number of brain areas including contralateral BA4, PMd, parietal lobe and ipsilateral cerebellum (Sharma, Jones, Carpenter, & Baron, 2008). Our previous work has demonstrated activation of the primary motor cortex (BA4) during motor imagery and execution, in the context of motor performance, is differentially affected by ageing (Sharma et al., 2008). We have additionally shown that that motor imagery and motor execution are increasingly bilateral within the motor cortex, and that imagery involves greater activation of BA4p than movement (Sharma & Baron, 2013) (Sharma et al., 2008). The broader question of the differential impact of ageing on cerebral networks for motor performance between motor imagery (MI) and motor execution (EM) has not been explored.

In order to probe whether there is an increased proportion of shared networks with age, we used tensor-independent component analysis (TICA), in a data-led approach. The major advantage of TICA is that it can demonstrate similarities between networks as well as differences. In this study, MI and EM are considered the same, so we are able to use a “blinded task” during the production of independent components (ICs). This means we make no assumptions about the presence or extent of overlap between MI and EM. We hypothesised that there would be shared components with EM and MI, areas which were recruited solely for EM and areas which were exclusive to MI. We also hypothesised that there would be age-dependent activation, (particularly in the sensory and somatosensory areas) as well as age independent activation (for example, in motor planning areas). The extent of shared networks demonstrates how age dependent common networks for EM and MI are.

## Methods

### Subjects

31 healthy volunteers were recruited through local advertisement. Mean age was 43.8 years (SD 17.1 Range 20–72), 13 were male. The data from subjects utilised here has also been used in our previously published work (Sharma & Baron, 2014). Subjects were medically screened and those recruited had no past or present medical history of any neurological, psychiatric or musculoskeletal disorders and were not taking regular medication. Analyses of data from these subjects has previously been undertaken (Sharma & Baron, 2014). All subjects were right handed as assessed by the Edinburgh Handedness scale (Oldfield, 1971). **Ethics Statement**. Informed consent was obtained from all individual participants included in the study, in accordance to the declaration of Helsinki. The protocol was approved by the Cambridge Regional Ethics Committee.

### Chaotic Motor Imagery Assessment Battery

All subjects were assessed using the Chaotic motor imagery assessment battery and were excluded if unable to perform motor imagery adequately. Chaotic motor imagery is defined as either an inability to perform motor imagery accurately or temporal uncoupling of otherwise accurate imagery (Sharma et al., 2006). Subjects were assessed using the Chaotic motor imagery Assessment Battery (Sharma et al., 2006) (Simmons, Sharma, Baron, & Pomeroy, 2008) (Heremans et al., 2011) (Sharma, Baron, & Rowe, 2009) (Sharma, Simmons, et al., 2009) which has three components that provide an objective measure of motor imagery compliance (see (Sharma et al., 2006) (Sharma, Baron, et al., 2009) (Sharma, Simmons, et al., 2009) for a full description). The first component used a hand mental rotation task to assess implicit motor imagery: subjects who score below 75% were excluded. The second component used variation of the fMRI task with a variable block length to ensure motor imagery performance. The third component used principles of motor control (more specifically Fitt’s Law, see (Sharma et al., 2006) (Simmons et al., 2008) (Heremans et al., 2011) (Sharma, Baron, et al., 2009) (Sharma, Simmons, et al., 2009)) to ensure subjects were using motor imagery: subjects using alternative cognitive strategies such as counting are excluded. During all motor imagery tasks, subjects were instructed to perform first-person motor imagery; not to view the scene from the 3rd person; and not to count, assign numbers or tones to each finger. Subjects were excluded if they were unable to perform motor imagery.

### Functional MRI

#### Motor paradigm and fMRI

As previously described (Sharma & Baron, 2015), The fMRI used an established block design (Oldfield, 1971) (Sharma, Simmons, et al., 2009) (Beckmann & Smith, 2005) with auditory pacing (1 Hz) of a right hand finger-thumb opposition sequence (2,3,4,5; 2…) with two separate runs (MI & rest and EM & rest). As activation patterns during MI may be unduly influenced if preceded immediately by execution (Lafleur et al., 2002), this was performed after MI. Subjects were instructed to keep their eyes closed. We used individually calibrated bilateral fibre-optic gloves (Fifth Dimension Technologies, SA) to monitor finger movements, exclude inappropriate movement and to assess the performance of motor imagery. After each motor imagery block, subjects confirmed whether the finger they were currently imagining was the correct ‘stop finger’ for the length of sequence. After scanning subjects were asked to rate the difficulty of motor imagery performance on a seven point scale (Alkadhi et al., 2005).

#### Data Acquisition

As previously described (Sharma & Baron, 2015), a 3-Tesla Brucker MRI scanner was used to acquire both T2-weighted and proton density anatomical images and T2*-weighted MRI transverse echo-planar images sensitive to the BOLD signal for fMRI (64664623; FOV 206206115; 23 slices 4 mm, TR = 1.5 s, TE 30 ms, Voxel Size 46464).

#### Image Analysis

Analysis was carried out using TICA (Beckmann & Smith, 2005), implemented in MELODIC (Multivariate Exploratory Linear Decomposition into Independent Components) Version 3.09, part of FSL (FMRIB’s Software Library, www.fmrib.ox.ac.uk/fsl), as previously described (Sharma & Baron, 2015). The first 12 volumes were discarded to allow for T1 equilibration effects. Preprocessing involved masking of non-brain voxels, voxel-wise de-meaning of the data, and normalization of the voxel-wise variance. Subject movement was less than 2 mm.

The preprocessed data were whitened and projected into a multidimensional subspace using probabilistic principal component analysis where the number of dimensions was estimated using the Laplace approximation to the Bayesian evidence of the model order (Smith et al., 2004). The whitened observations were decomposed into sets of vectors which describe signal variation across the temporal domain (time courses), the session/subject domain, and the spatial domain (maps) by optimizing for non-Gaussian spatial source distributions using a fixed-point iteration technique (Hyvärinen, 1999). Estimated component maps were divided by the standard deviation of the residual noise and thresholded by fitting a mixture model to the histogram of intensity values. The time course of each IC was then entered into a general linear model of the convolved block design of Task versus Rest.

An IC was considered to be involved in MI or EM if a one-way t-test found the subject scores to be significantly different from zero across subjects, and there was plausible anatomical localization and normalized response. When an IC was significantly involved in both tasks, then a paired t-test (p < 0.05 corrected for multiple comparisons) was performed on the subject score for each task. For all subjects, scores for MI and EM of each remaining component were correlated with subject age (Spearman p < 0.05). Due to the degree of smoothing, we did not distinguish between BA4a and BA4p.

## Results

### Behavioural Results

As previously reported (Sharma & Baron, 2014), 5 subjects were excluded; three because of a failure to perform implicit motor imagery satisfactorily (mean Score 69.9%); two because of the use of alternative cognitive strategies (mean break point +23%). The remaining 26 subjects (12 male) performed adequately on all aspects of the hand rotation task (mean 94.4% SD 4.3%) fMRI simulation and Fitts’ law (mean break point 17.4% less for motor imagery then movement) as well as during the MRI session. Median post MRI motor imagery scores (MIS) was Right hand = 6. No subject failed to either suppress movement or showed evidence of non-compliance during the fMRI paradigm.

### fMRI Data

No distinction was made between tasks until the final stage of processing. As 26 subjects performed two tasks, MI and EM, 52 “blinded” tasks were processed. As no distinction was made between imagery and EM during the generation of the ICs, we use the term “blinded.” A subject score for each IC was produced that included the effect size for the 52 blinded tasks for the associated spatiotemporal process shown in the spatial map.

Thirty-seven ICs were defined by TICA. ICs that identified artifact recognized by previously published patterns and high frequency were excluded by visual inspection. ICs driven by outliers or not significant across either task were also excluded. Only ICs that were driven by activity during activation blocks were included in analysis. Therefore, only components in which the subject scores were significantly different from zero (for either motor imagery or execution) were included.

Eight ICs remained. Table 1 summarizes the areas involved, labelled using the SPM Anatomy toolbox (Eickhoff et al., 2005) (Eickhoff et al., 2007) (Eickhoff, Heim, Zilles, & Amunts, 2006) and the Talairach Atlas (Lacadie, Fulbright, Rajeevan, Constable, & Papademetris, 2008) after MNI conversion (Lacadie et al., 2008).

**Table 1.**
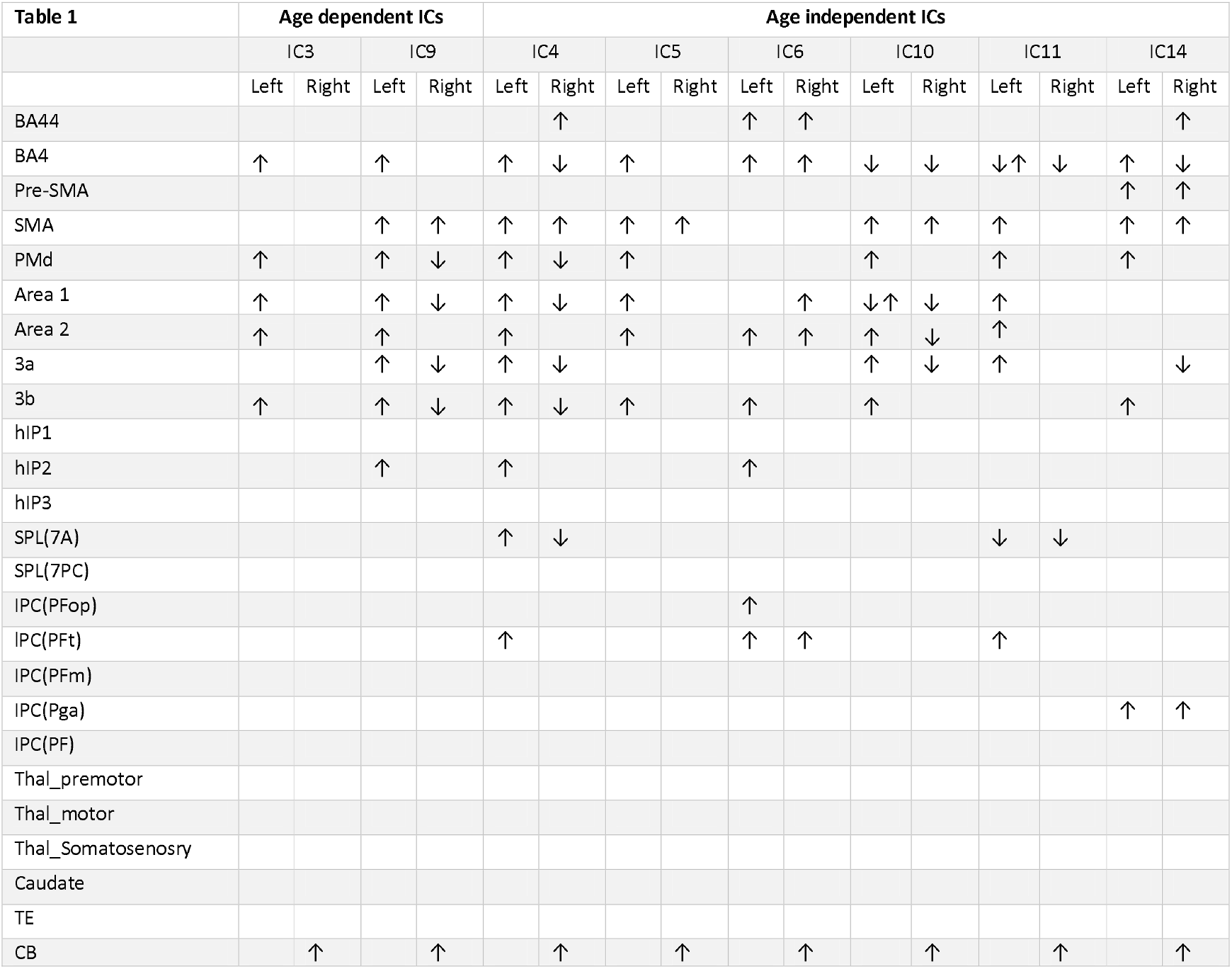
Regions activated (→) or deactivated (↓) within each independent component. All components are shared between motor imagery (MI) and motor execution (EM). Changes in activation ICs 3 and 9 were dependent of age.

### Independent Components

Figures 1 and 2 show the whole brain activations and deactivations, the time course, subject scores, and percentage of total variance explained.

**Figure 1.**
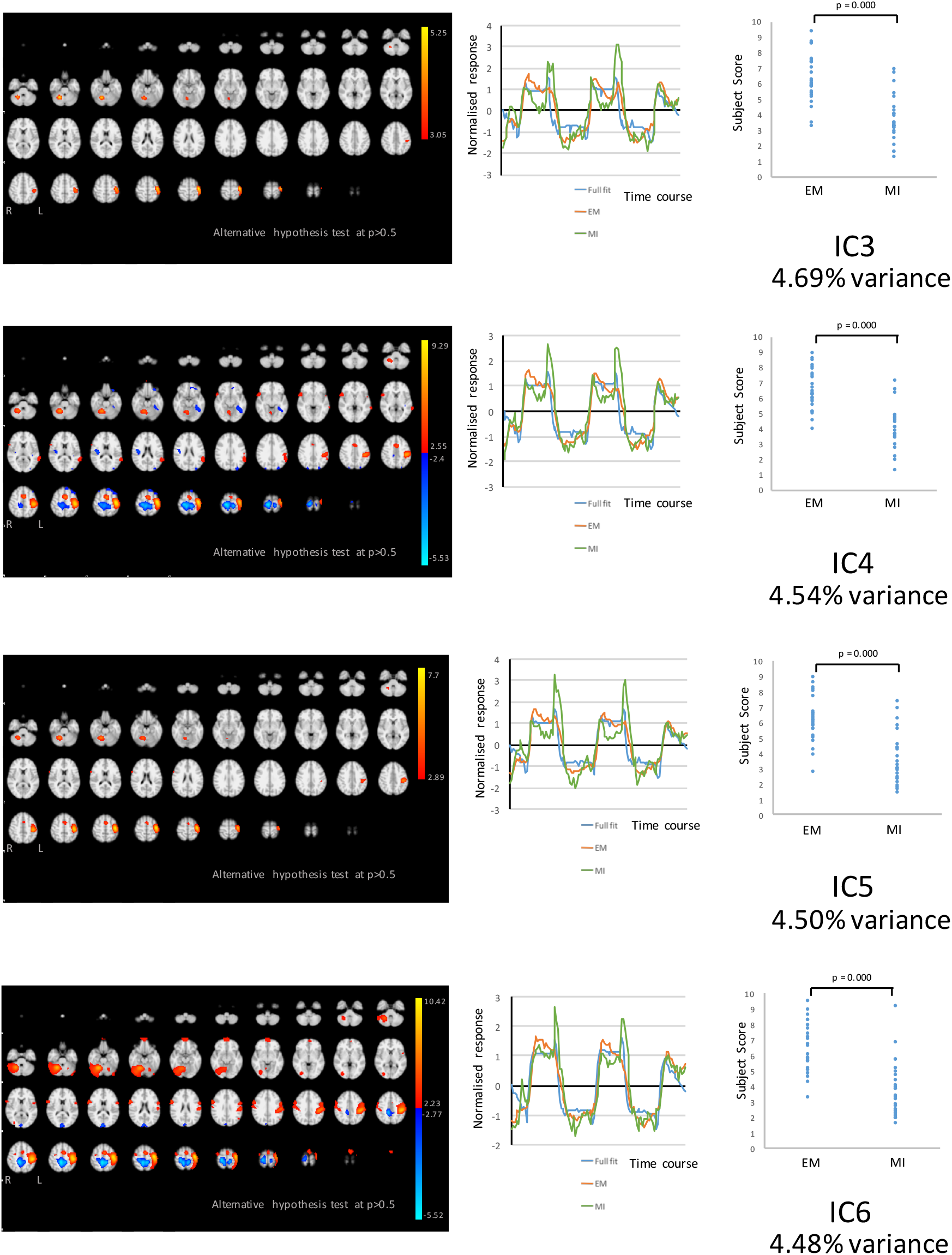
The figures show the involvement of each IC across the whole brain with a standard threshold of p > 0.5 (alternative hypothesis test) and the variance it accounts for out of the total explained variance. Only ICs that survived all stringency tests detailed in methods are shown in Figure 1 and Figure 2. The scales show the transformed z-score, orange is activation, and blue is deactivation. The normalized time course response is shown for each task and the full model fit (full model fit = blue, EM = red, MI = green). The subject scores for MI and EM are shown for each component.

**Figure 2.**
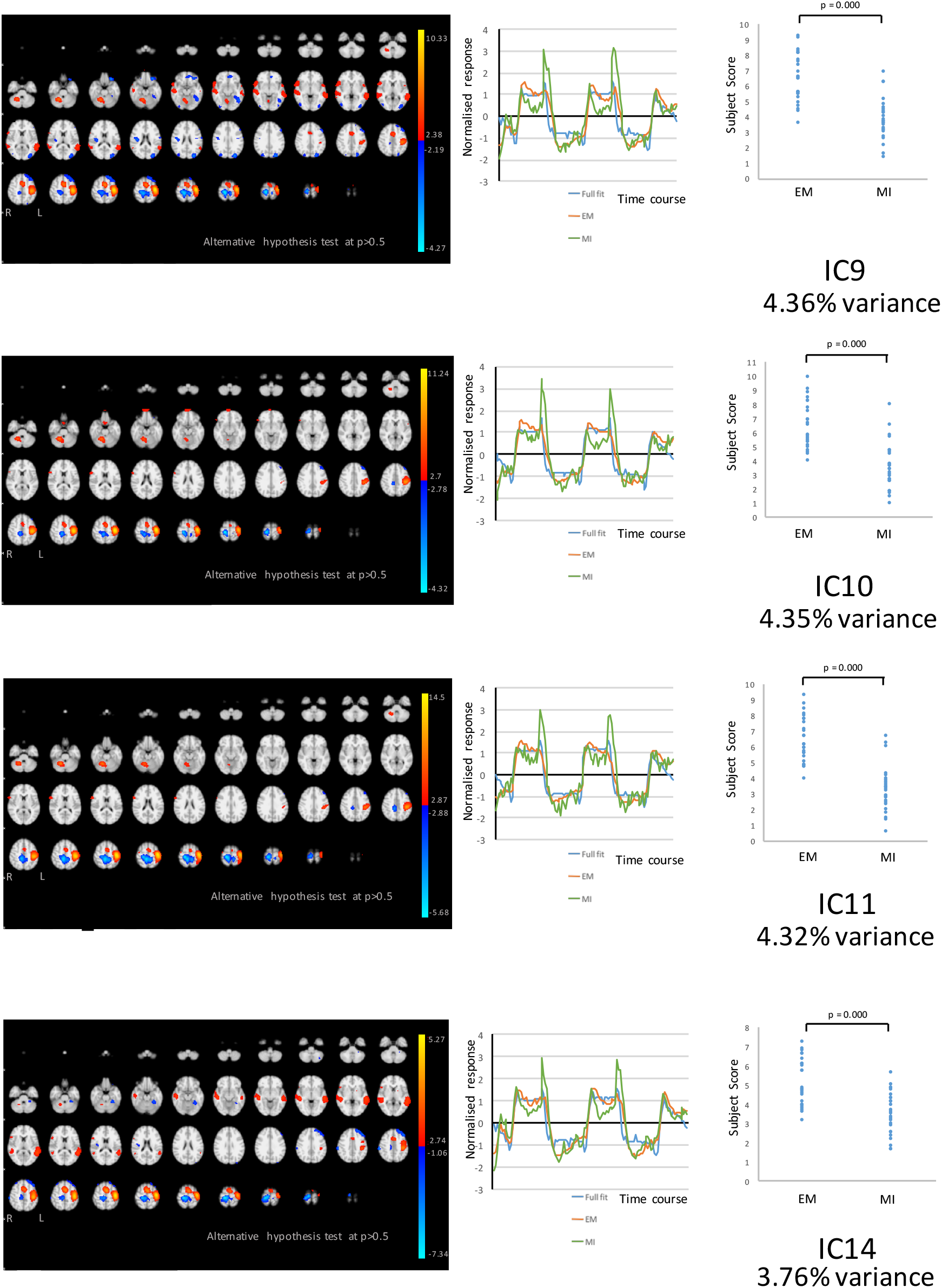
The figures show the involvement of each IC across the whole brain with a standard threshold of p > 0.5 (alternative hypothesis test) and the variance it accounts for out of the total explained variance. Only ICs that survived all stringency tests detailed in methods are shown in Figure 1 and Figure 2. The scales show the transformed z-score, orange is activation, and blue is deactivation. The normalized time course response is shown for each task and the full model fit (full model fit = blue, EM = red, MI = green). The subject scores for MI and EM are shown for each component.

**Figure 3.**
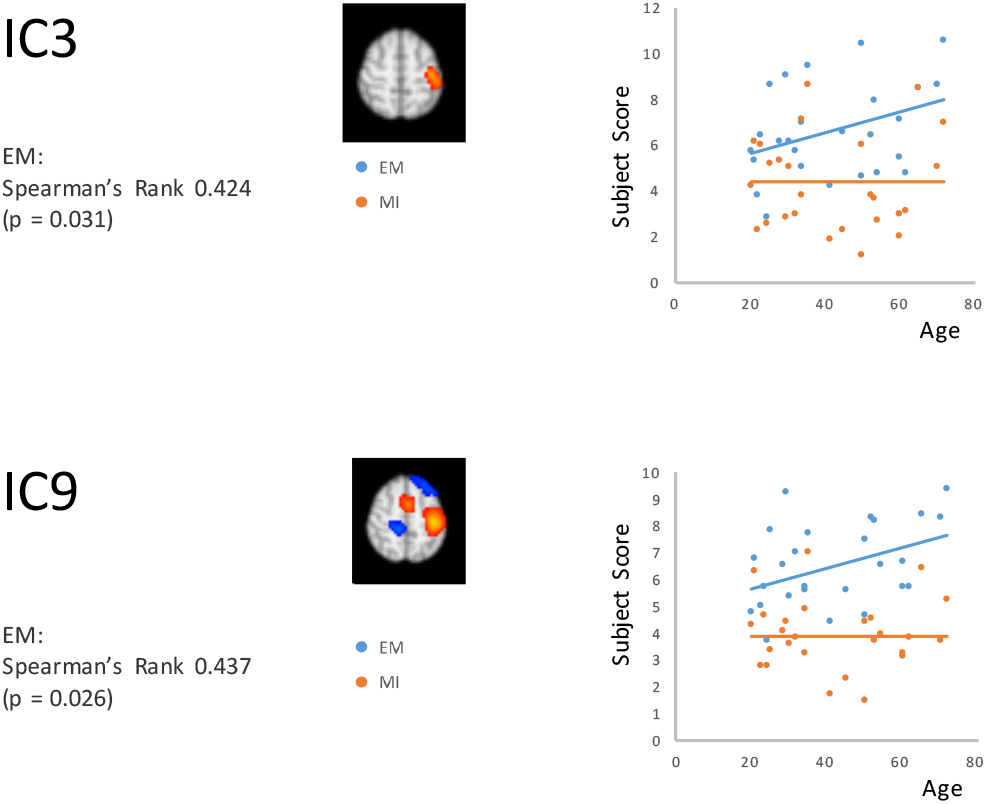
Only IC3 and IC9 have statistically significant correlations between EM and MI with age. Spearman’s Rank correlations are shown.

All ICs were similar in variance, with a range between 3.76% and 4.69%. The total variance explained was 35%. All ICs were shared between EM and MI (subject scores significantly greater than zero for both tasks in both groups), with activation in EM being greater than MI. There were no ICs where EM was equal to MI or MI-only ICs. There was no relationship between MIS and ICs.

Activation of the contralateral BA4 and ipsilateral cerebellum was consistent across the components. This further supports the known overlap between EM and MI. Bilateral SMA activation was prominent in several components (IC4, IC5, IC9, IC10, IC11(Left) & IC14). Parietal lobe activation was present in IC4, IC6, IC11 and IC14.

A number of IC’s showed deactivation of the ipsilateral BA4 (IC 4,10,11 & 14), sensory cortex (IC9,IC4)and secondary motor areas (IC4, IC9).

### Correlation with Age

ICs 3 and 9 positively correlated with age for EM (IC3 rho = 0.38, p = 0.033; and IC9 rho = 0.39, p = 0.026) see **Figure 2**. There was no correlation MI for these components.

These components explained 9.1% of the variance. Subjects scores of these components increased with greater age for EM only. IC3 is notable in contralateral activation of BA4, sensory cortex, PMd and ipsilateral cerebellum. There was no activation (or deactivation) of ipsilateral motor areas or SMA. Whereas IC9 activated similar areas, but also included SMA and deactivation of the ipsilateral motor and sensory cortex.

## Discussion

This work furthers knowledge on the overlap between MI and EM and extends the work regarding the effects of age. For age-related components, activation for EM and not MI appeared to positively relate to components including the primary motor cortex, sensory cortex and cerebellum. The overlap in EM and MI over age-dependent components with evidence of age related changes in EM only highlights the importance of ageing and supports the differences use of MI as alternative means to access the motor system.

There were four key findings. First, eight ICs exhibited a pattern consistent with activation during MI and EM. Second, these ICs accounted for 35% of variance. Third, there was bilateral activation across cortical areas and the cerebellum. Fourth, two ICs had statistically significant correlations between EM (but not MI) with age are reported: components 3 and 9. The other six ICs were age independent. Fourth, there were only EM and MI shared components, with no MI-only components, which was not entirely unexpected. We explore this below.

Given six components are age independent and activated in EM > MI, it is likely that cortical and subcortical motor networks are in large part maintained over life, and that differences in performance are likely to be attributable in large part to distal changes.

In this study, all ICs are involved in MI and EM. It is biologically plausible that motor imagery and execution share common networks as part of precognition of a movement. Experimental work using this cohort has demonstrated formally that there are shared networks for MI and EM (Sharma & Baron, 2013). Similar to this study, there were no MI only components. However, in contrast to previous findings, there were no EM-only components. In previous work (Sharma & Baron, 2013), EM only components have involved bilateral activation of primary and secondary motor cortical areas and the cerebellum. It is possible that differences with age were too subtle for this sample size. It is unlikely that subclinical extension that was not detected, given the use of data from these subjects in previous studies. Regarding age dependent components, it is possible that changes with age are more evident in shared networks.

It is more difficult to interpret reasons for lack of EM-only components. Secondary motor and subcortical areas, which have featured in EM-only components in previous work (Sharma & Baron, 2013), are included in areas of activation of these components. This suggests that despite our attempts to prevent movement, there may have been discharge via the CST. However, against this is that age only related to EM rather than both conditions. A study examining using fMRI to probe changes in activation of finger movements with ageing demonstrated a divergence of MI and EM with age. The differences in MI related to the extent of activation, but in EM, elderly subjects showed additional activations in the left inferior occipital gyrus, in the right paracentral lobule, and in the right pre-SMA (Zapparoli et al., 2013). Previous work using fMRI in young and old subjects in motor tasks of differential complexity (dictated by extent of isodirectionality) have shown that in old subjects, accessory motor areas are recruited to a greater differential extent in more complex motor task, so it may be that the finger tapping task used in this study was insufficiently complex to demonstrate accessory motor area recruitment (Heuninckx, Wenderoth, Debaere, Peeters, & Swinnen, 2005). An additional factor to consider is task performance of old subjects relative to younger subjects. Previous work has demonstrated an inverse correlation between extent of access motor network recruitment and performance in a button press motor task (Mattay et al., 2002), so it may be the case that strict selection criteria that were used in this study has led to a selection bias away from subjects that would demonstrate accessory motor system recruitment.

In this study, all ICs had activation greater in EM than in MI. This is in contrast to previous studies which have also produced ICs where MI has equal or greater activation versus EM (Sharma & Baron, 2013) (Sharma & Baron, 2015). There are four possible reasons for this. First, all ICs may involve a distal component where there is corticospinal tract activation (Sharma & Cohen, 2012) and sensory feedback to the motor cortex of the movement itself (Christensen et al., 2007) (Cullen, 2004). Second, MI components may be present and suppressed. Against this would be the lack of existence of EM only or MI only components.

In favour of this is evidence of corticospinal suppression with motor imagery of movements induced by transcranial magnetic stimulation (Sohn, Dang, & Hallett, 2003). Third, motor imagery may itself have an excitatory effect distally to the cortex, within the corticospinal tract. Electromyographic responses of very short latencies with motor imagery, similar to that elicited by reflexes, provide evidence in favour of this (Li, Kamper, Stevens, & Rymer, 2004). Finally, there may be a mechanism of stopping movement that this block design does not pick up. In terms of energy expended, not moving has a similar cost to moving (Noorani & Carpenter, 2017). Movement cancellation is determined by a network including the basal ganglia and frontal areas (Noorani, 2017). Impending tasks are required to explore cancellation, such as the Go/No-Go paradigm (Noorani, 2017). An event related design would therefore be a useful complement to this experiment.

There were two ICs where EM related to age. Age dependent bilateral changes have been predicted from previous predictions on changes to activation with ageing, such as the Hemispheric Asymmetry Reduction in Older Adults (HAROLD) model (Cabeza, Anderson, Locantore, & McIntosh, 2002). Primary motor cortex activity during a finger thumb opposition task has been shown as increasingly bilateral with age (Naccarato et al., 2006).

IC3 and IC9 represent changes in activation in a number of biologically plausible areas. For IC3, BA6 has roles in motor sequencing (Bischoff-Grethe, Goedert, Willingham, & Grafton, 2004), motor learning (Honda et al., 1998), motor imagery (Malouin, Richards, Jackson, Dumas, & Doyon, 2003), and coordination between limbs (Ehrsson et al., 2000); BA10 may have roles in reward processing and decision making (Rogers et al., 1999) or recall of event based memories as part of imagery in this context (Okuda et al., 2007); BA46: may be utilized for retrieval or mirror neuron activation (Buccino et al., 2004) (Dolan, Lane, Chua, & Fletcher, 2000). For IC9, activation of BA3b relates to finger proprioception (Carey, Abbott, Egan, & Donnan, 2008), and changes in activation of BA30 and B31 may be driven by their roles in motor learning (Tracy et al., 2003) or integration of sensory and motor processes (Schubert, von Cramon, Niendorf, Pollmann, & Bublak, 1998).

These age-dependent components likely arise due to two types of age-dependent change that occur in circuits involved with EM. There are two explanations for these changes. First, motor execution takes increasing advantage of areas activated in motor imagery, and there is increasing prominence of shared circuits with age, hence MI activation will remain similar, but EM activation may change with age. Second, there is also activation of distal motor networks in motor execution. This increase in activation supports a move from automatic motor control to motor control requiring more input from motor imagery with age. Increasingly bilateral motor cortex activation is correlated with contralateral ventral premotor cortex activation. Studies have found increased multisensory activation including the somatosensory cortices (Zwergal et al., 2012) visual cortical areas (Zwergal et al., 2012) (Nedelko et al., 2010) and the superior parietal lobule (Nedelko et al., 2010).

## Conclusion

This study demonstrates the prominence of shared cerebral networks between MI and EM. Comparing MI and EM with ageing using a tensor independent component analysis, the majority of variance is age independent. There are EM related changes with age, but no changes in activation for MI.

Age dependent ICs include a diffuse network of brain areas, spanning sensorimotor and cerebellar changes. Age dependent variance includes bilateral changes to activation. The predominance of age-independent networks over age-dependent networks strengthens the rationale for using MI to access the motor networks for therapeutic purposes in patients with neurological disease, irrespective of age.

## Ethics

This study was conducted with the human subjects’ understanding and consent. The Cambridge Regional Ethics Committee approved the protocol.

## Acknowledgements: Funding

This work was supported by The Stroke Association (TSA 2003/10) and the Medical Research Council (MRC G0001219). N Sharma receives a proportion of funding from the Department of Health’s National Institute for Health Research (NIHR) Biomedical ResearchCentres funding scheme for UCLH.

